# Data-independent acquisition parallel accumulation-serial fragmentation (diaPASEF) analysis of the separated zebrafish lens improves identifications

**DOI:** 10.1101/2025.03.13.642445

**Authors:** Sarah R. Zelle, W. Hayes McDonald, Kristie L. Rose, Hassane S. Mchaourab, Kevin L. Schey

## Abstract

Ocular lens fiber cells degrade their organelles during differentiation to prevent light scattering. Organelle degradation occurs continuously throughout an individual’s lifespan, creating a spatial gradient of young cortical fiber cells in the lens periphery to older nuclear fiber cells in the center of the lens. Therefore, separation of cortical and nuclear regions enables examination of protein aging. Previously, the human lens cortex and nucleus have been studied using data-independent acquisition (DIA) proteomics, allowing for the identification of low-abundance protein groups. In this study, we employed data-independent acquisition parallel accumulation-serial fragmentation (diaPASEF) proteomics on a timsTOF HT instrument to study the zebrafish lens proteome and compared results to a standard DIA method employed on an Orbitrap Exploris 480. Using the additional ion mobility gas phase separation of diaPASEF, peptide and protein group identifications increased by over 200% relative to an orbitrap DIA method in the zebrafish lens. With diaPASEF, we identified 13,721 and 11,996 unique peptides in the zebrafish lens cortex and nucleus, respectively, which correspond to 1,537 and 1,389 protein groups. Thus, separation of the zebrafish lens into cortical and nuclear regions followed by diaPASEF analysis produced the most comprehensive zebrafish lens proteomic dataset to date.

## Introduction

The lens is a transparent organ that focuses light onto the retina. With age, the lens becomes opacified and scatters light, a condition known as cataract, the leading cause of blindness worldwide.^1–3^ Cellular organelles act as potential sources of light scattering, so during lens fiber cell differentiation, organelles are degraded.^1^ As we age, continual lens fiber cell differentiation creates a spatial gradient of young cortical fiber cells to old nuclear fiber cells (Figure 1).^2–4^ Due to the loss of organelles during differentiation, there is no protein turnover in nuclear lens fiber cells, and proteins synthesized at birth remain in the nucleus for the lifespan of an individual. ^5^ Lens fiber cell differentiation thus creates a continuum of regions with distinct proteomes in the lens, necessitating the separation of lens regions to examine age-related proteome changes. Conventionally, the lens is separated into organelle-containing cortical and organelle-free nuclear regions for spatiotemporal proteomic analyses.^5,6^

**Figure 1.**
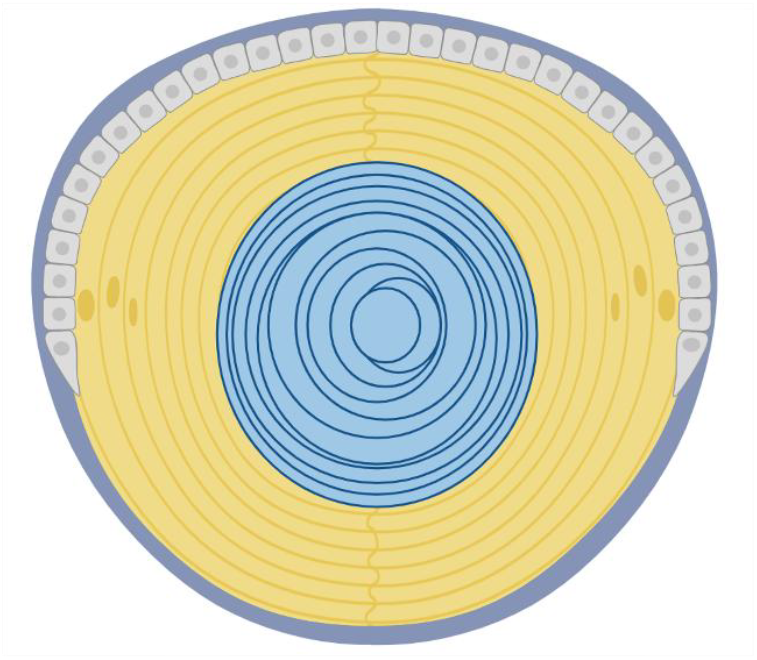
Diagram of the zebrafish lens. Epithelial cells (light gray) differentiate into lens fiber cells. The outer layer of lens fiber cells is the cortex (yellow), while the inner layer is the nucleus (blue). The lens is enclosed in the capsule, which is comprised of basement membrane (dark gray). Figure created using Biorender.com.

Abundance-based data-dependent acquisition (DDA) mass spectrometry proteomics and labeling methods such as tandem mass tags have been used to quantify proteins and post-translational modifications in the lens.^7–12^ However, due to the high concentration of crystallins, up to 300 mg/mL in the human lens, these methods are challenged to identify and quantify low-abundance proteins.^13^ Thus, new data-independent acquisition (DIA) mass spectrometry proteomics methods were developed to study the human lens proteome.^14,15^ Using a 95-minute liquid chromatography gradient combined with a library-free DIA method on an Orbitrap Exploris 480 instrument to interrogate the human lens proteome, Cantrell and Schey identified 39,292 peptides, corresponding to 4,476 protein groups in the lens cortex. In contrast, a DDA method on the same nstrument identified 9,448 peptides and 1,739 corresponding protein groups in the lens cortex. These results show that by using DIA there was an almost 250% increase in peptide identifications and an almost 400% increase in protein group identifications. Fewer peptides and protein groups were identified in the lens nucleus region relative to the cortex, but the DIA method also identified more peptides and protein groups than the DDA method. diaPASEF (data independent acquisition parallel accumulation serial fragmentation) mass spectrometry proteomics, takes advantage of an additional ion mobility gas phase separation device and high-speed mass analysis to increase the number of peptide and protein group identifications.^16^ Here, we pioneer the use of diaPASEF mass spectrometry proteomics to study the zebrafish lens and compare our results to data acquired using the previously reported orbitrap DIA method on the same tissue.

Previously, limited zebrafish lens proteomic analyses have been published, focusing either on development or quantifying changes in α-crystallin abundance using DDA mass spectrometry proteomics.^17–20^ Zebrafish have several attributes that make them ideal model organisms for studying the lens proteome. For instance, zebrafish undergo a similar lens organogenesis process and have an analogous lens structure to humans.^21,22^ However, the zebrafish lens has traditionally been analyzed intact due to its small size. Thus, for the first time, the nuclear and cortical regions of the zebrafish lens were separated for proteomic analysis, allowing for the examination of lens fiber cell aging. By utilizing diaPASEF proteomics, we collected the most comprehensive proteomic dataset of the zebrafish lens to date.

## Materials and Methods

All reagents were purchased from Sigma-Aldrich (St. Louis, MO) and S-Trap micro columns were acquired from Protifi (Farmingdale, NY). Mass spectrometry grade trypsin and other remaining materials were purchased from Fisher Scientific (Waltham, MA) unless noted otherwise.

### Zebrafish Maintenance and Breeding

AB-wild type strain zebrafish (Danio rerio) embryos were raised to 10 months of age. Zebrafish were kept in 30 mg/L instant ocean in deionized water at 28.5 °C and on a 14:10 hour light/dark cycle. The Vanderbilt University Institutional Animal Care and Use Committee authorized all zebrafish procedures.

### Dissection and Separation of the Lens

Fresh lenses from three biological replicates were dissected from the eye and placed in cold PBS.^23^ To separate the lens cortical and nuclear regions, 5 μL of cold 100 mM Tris pH 7.8, 1.5 mM ethylenediaminetetraacetic acid (EDTA) buffer was first placed in the neck of Biotik xPY4 20 μL pipette tips. Lenses were gently dried on a Kimwipe using tweezers and put into the buffer droplet in the pipette tip. The tips were spun at 5,000g for 30 seconds at 4 ºC in centrifuge tip adapters inserted into microcentrifuge tubes. The centrifugation process shears off the cortical region of the lens into the eluted solution while the lens nucleus remains intact and stuck in the tip. The tips were then cut using a razor blade at a point just below where the lens nucleus was held. Tweezers were used to free the nucleus by pushing it towards the larger end of the tips and spinning again to elute the intact nucleus into the buffer containing the separated cortex region. To wash the tips of any lingering cortical tissue, 10 μL of cold buffer was added and the tips were spun again. The buffer containing the cortical region was then transferred to another tube, leaving the intact nucleus inside the initial tube. The exterior of the nucleus was then washed by suspending it in 10 μL of cold buffer to remove any remaining attached cortical fiber cells. The wash buffer was then transferred to the cortical fraction and the separated regions were then snap-frozen for storage at -80 ºC.

### Preparation of samples for mass spectrometry proteomic analysis

Separated tissue samples were thawed and 50 μL of 100 mM Tris pH 7.8, 1.5 mM EDTA, 5% sodium dodecyl sulfate (SDS) buffer was added to the nuclear samples, while 25 μL of 100 mM Tris pH 7.8, 1.5 mM EDTA, 10% SDS buffer was added to the cortical samples. All samples were homogenized using an electric pestle and spun down at 18,213g for 5 minutes to pellet tissue. Protein concentration in the supernatant was measured using a bicinchoninic assay. Cysteines in 10 μg of protein were reduced for 1 hour at 56 °C using 10 mM of dithiothreitol and alkylated for 30 minutes in the dark at room temperature using 50 mM of iodoacetamide. Proteins were acidified by adding a 1:10 ratio of 27.5% phosphoric acid and then precipitated onto an S-Trap micro spin column by adding a 6:1 ratio of 100 mM triethylammonium bicarbonate (TEAB) in 90% methanol to sample and centrifuged at 4,000g for 30 seconds. Samples were washed three times with 50% chloroform/50% methanol, followed by four additional washes with 100 mM TEAB in 90% methanol. Proteins were digested by adding 50 mM TEAB with a 1:10 ratio of 10 μg sample to 1 μg of mass spectrometry grade trypsin and incubated for 2 hours at 47 °C. Digested peptides were eluted from S-trap sequentially by adding 50 mM TEAB, 0.2% formic acid (FA), 50% acetonitrile, and 0.2% FA solutions and centrifuging at 4,000g for 1 minute each. Eluted peptides were dried with a vacuum concentrator and stored at -80 ºC.

### Data Acquisition

Peptides were resuspended in 0.2% FA to a concentration of 200 ng/μL. Peptide concentrations were estimated using a Nanodrop 2000 Spectrophotometer (Thermo Scientific), and samples were adjusted to 50 ng/μL. 0.015% n-dodecyl-ß-D-maltoside was added to diaPASEF samples. diaPASEF data were acquired on 150 ng of peptides using a 30-minute aqueous to organic gradient method delivered via a nanoELUTE2 on a PepSep column with 75-micron internal diameter, 25 cm length, and 1.5-micron particle size coupled to a timsTOF HT instrument (Bruker) using a 20-micron Captive Spray emitter. diaPASEF data were collected in 12 PASEF ramps from 0.75 to 1.3 1/k_0_ covering 350 to 1250 m/z via variable windows ranging in size from 12.27-122.81 Th with 50 ms accumulation time. Orbitrap DIA data were acquired on 150 ng of peptides using a 95-minute aqueous to organic gradient method delivered via a Dionex Ultimate 3000 UHPLC on a 360 μm outer diameter × 100 μm internal diameter column packed with 20 cm of 3 °m C18 reverse phase material (Jupiter, Phenomenex) coupled to an Orbitrap Exploris 480 instrument (Thermo Scientific). Precursor spectra from 385-1025 murs continuously throughout an indivi windows of DIA MS/MS spectra from 400-1000 m/z or thirty-one 20 m/z windows of DIA MS/MS spectra from 390-1010 m/z.^14,15^

### Data Analysis and Visualization

Database searching for all files was performed using DIA-NN 1.9 with the default library-free settings and the reannotate and contaminants options selected.^24,25^ A UniProt Swiss-Prot and TrEMBL zebrafish FASTA database (UP000000437, downloaded 9/10/2024, 25,999 entries with only one protein per gene) was used. Match between runs was turned on and MS1 and MS2 mass accuracy were fixed at 10 ppm. The default fixed N-terminal methionine excision and cysteine carbamidomethylation modifications and no variable modifications were selected. Searching was performed on an Intel Core i7-10700 CPU at 2.90 GHz.

Data were analyzed and visualized using custom R scripts on non-contaminant peptides having < 1% q-value and at least 2 unique peptides per protein group. DIA-NN normalization was rejected, and median normalization was performed separately on files for each instrument (Supplemental Figure 1).^14,15^ Abundances were calculated using the diann_maxlfq function from the diann R package.^24^ Figures were generated in R using ggplot2, ggvenn, and EnhancedVolcano packages. The prcomp package was used to perform principal component analysis (PCA) on un-transformed for all protein groups without missing values in the diaPASEF and orbitrap DIA datasets separately. A Welch’s t-test between cortical and nuclear samples was performed on all proteins. Results were plotted using a volcano plot with significance cutoffs at 0.05 *p* value and +/-1.5 log_2_ fold change. PANTHER was used for overrepresentation analyses of all proteins with a significant *p* value < 0.05 as determined via Welch’s t-test using the 06/17/2024 GO database.^26^ Statistically significant cortical and nuclear proteins from the diaPASEF and orbitrap DIA datasets were analyzed separately using a Fisher’s exact test with false discovery rate (FDR) calculated. All identified proteins from both data acquisitions comprised the reference proteome, and only parent GO terms with FDR *p* value < 0.05 were considered. Data are available via ProteomeXchange with identifier PXD059560.

## Results and Discussion

Due to the additional gas phase separation and the higher instrument acquisition speed, the diaPASEF method was predicted to increase the number of peptide and corresponding protein group identifications relative to orbitrap DIA in the zebrafish lens cortex and nucleus. Results revealed that among three biological replicates an average of 3,584 unique peptides were identified using an orbitrap DIA method in the zebrafish cortex and an average of 3,103 unique peptides were identified in the zebrafish nucleus. However, by using diaPASEF these numbers increased to an average of 13,721 unique peptides identified in the zebrafish lens cortex and 11,996 unique peptides in the nucleus, representing about a 260% average increase in peptide identifications in both regions (Figure 2A). In the orbitrap DIA dataset, these peptides can be assigned, on average, to 511 unique protein groups in the cortex, and 432 unique protein groups in the nucleus. In the diaPASEF dataset, these numbers increased to an average of 1,537 unique protein groups in the cortex and an average of 1,389 unique protein groups in the nucleus (Figure 2B). These identifications represent about a 330% increase in cortical protein group identifications and about a 310% increase in nuclear protein group identifications using diaPASEF.

**Figure 2.**
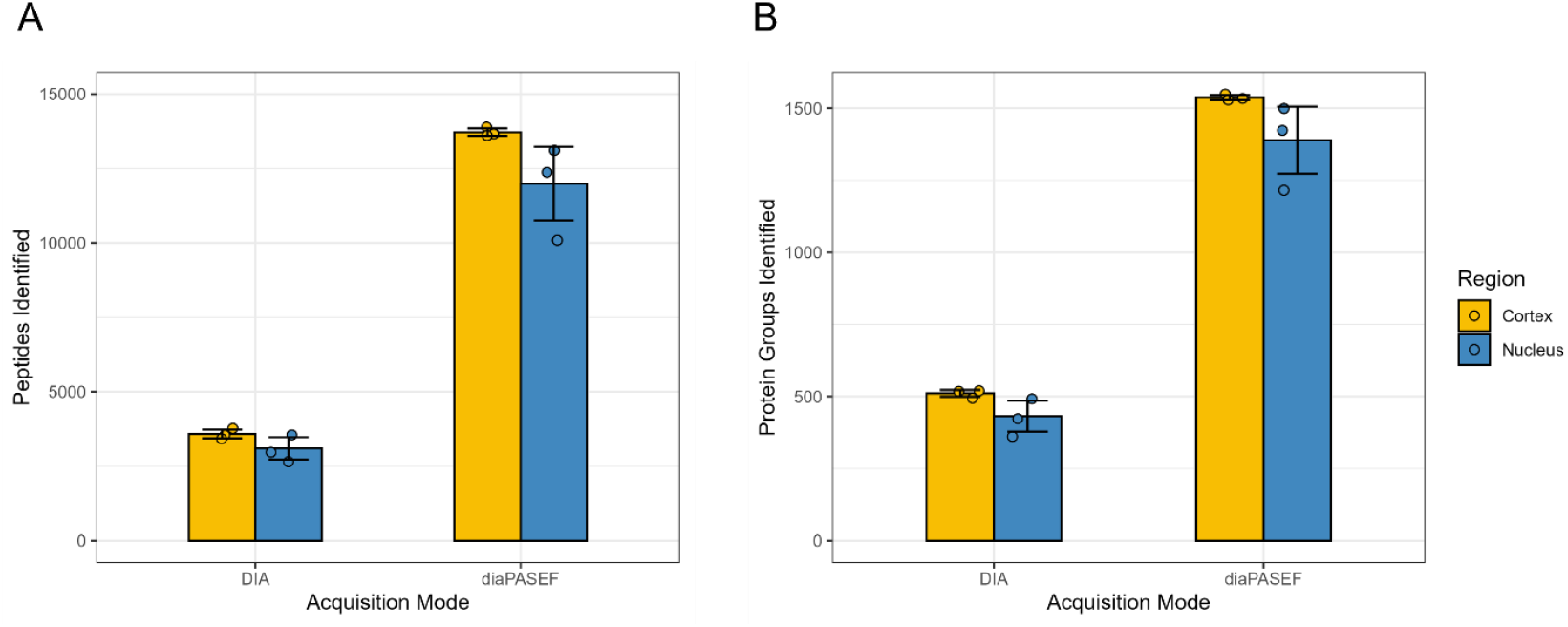
Bar plots of orbitrap DIA and diaPASEF results for three biological replicates displayed by acquisition mode and region. **A)** number of peptides identified and **B)** number of corresponding protein groups identified. A total of 16,718 unique peptides and 1,674 protein groups were identified throughout all sample types and data acquisition strategies.

We next compared the overlap between identifications on the peptide and protein group level in the zebrafish cortex and nucleus for both data acquisition methods. On the peptide level, about 20% of all identifications in the lens cortex and nucleus were detected using both orbitrap DIA and diaPASEF. However, over 70% of all identified peptides were found exclusively in the diaPASEF datasets in both regions, suggesting ion mobility is needed to identify these additional peptides (Figure 3A) within a faster acquisition time. At the protein group level, there was more overlap, with more than 30% of all protein groups identified in both regions using both acquisition methods. Over 60% of all protein groups were found only in the diaPASEF datasets for both regions, suggesting that diaPASEF gives a more complete protein coverage in the zebrafish lens (Figure 3B). The diaPASEF datasets also had, compared to the orbitrap DIA datasets, a lower median MS2-based peptide intensity percent coefficient of variation (% CV) and proportion of peptides with a lower % CV (Supplemental Figure 2A and 2B).

**Figure 3.**
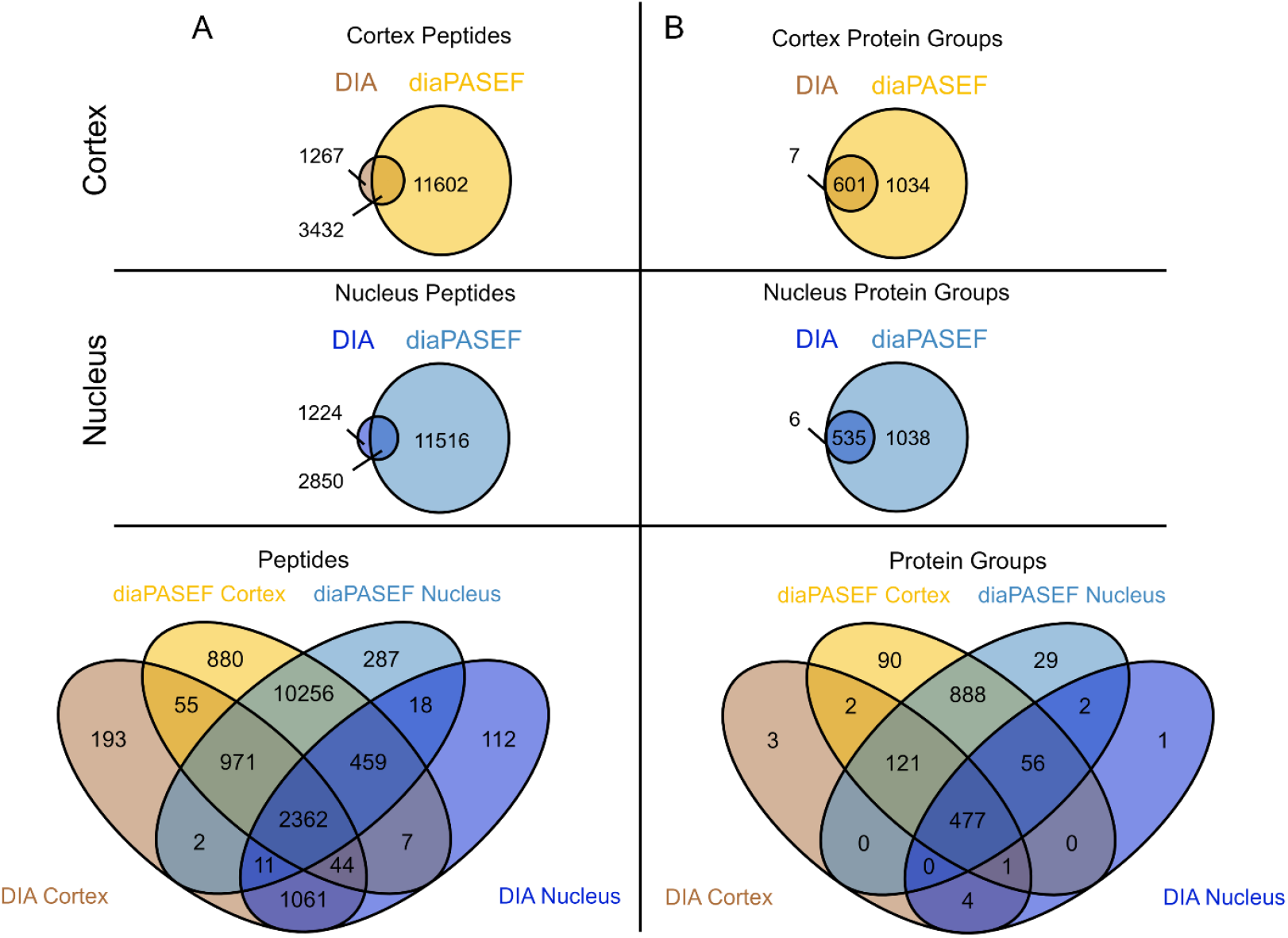
Venn diagrams showing the overlap between acquisition methods for. **A)** peptides and **B)** corresponding protein groups identified in the zebrafish lens cortex and nucleus. All peptides and protein groups found in at least one biological replicate were considered, with duplicates removed.

Additionally, there is a notable overlap between the cortex and nucleus regions in identifications on both the peptide and protein group levels in both data acquisition methods (Supplemental Figure 3A and 3B). Differences in peptide and protein group identifications were likely due to protein modification, truncation, degradation, or differences in protein synthesis pathways, processes associated with nuclear fiber cell protein aging.^5,11,27,28^ A high degree of similarity was also found between replicates (Supplemental Figure 4). However, unique peptides and protein groups found in only one replicate are likely a result of low abundance peptides being less reproducible and the high biological variance found in zebrafish.^29–31^ These results demonstrate that diaPASEF facilitates the quantification of more unique peptides and protein groups in the zebrafish lens compared to the orbitrap DIA method, allowing for deeper investigation into lens biology.

**Figure 4.**
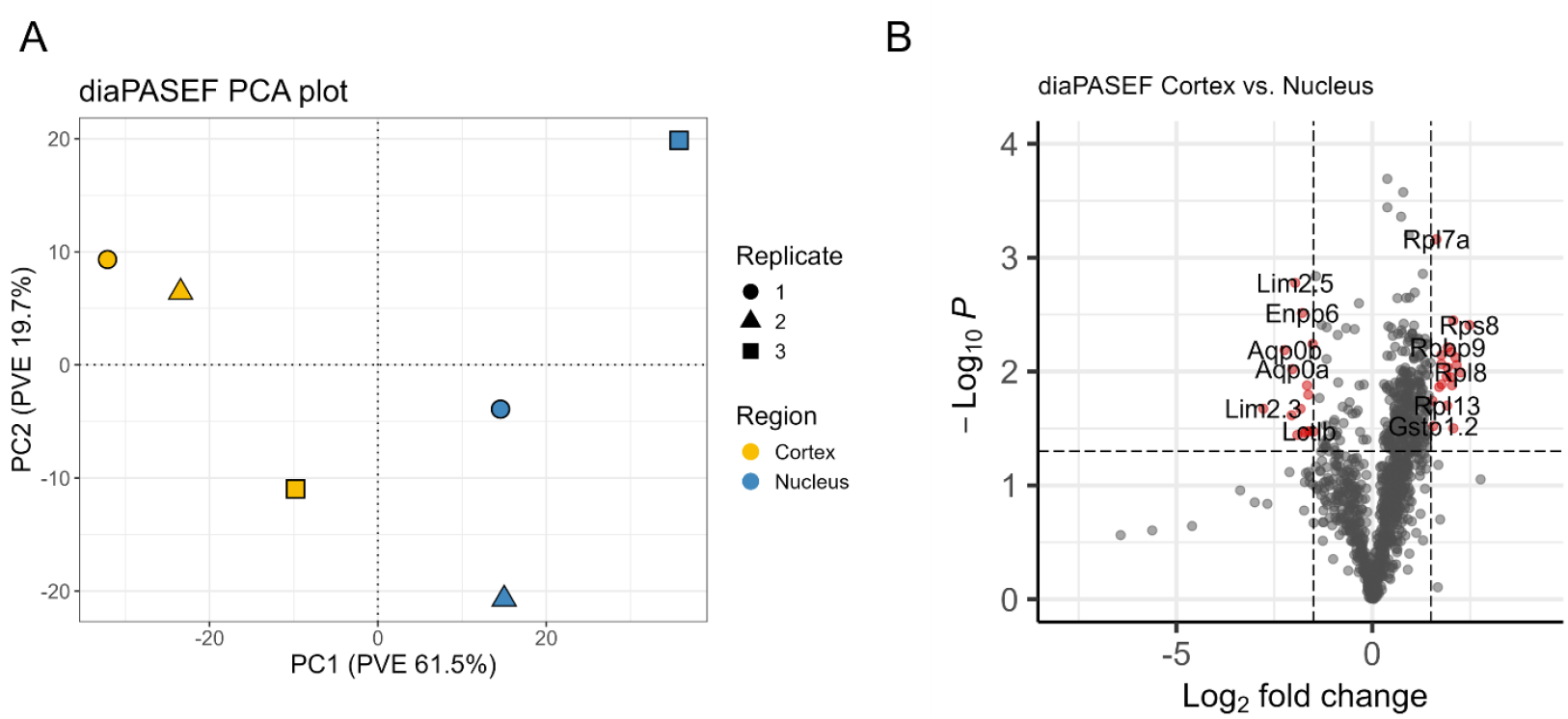
Analysis of diaPASEF data. **A)** PCA plot separated by replicate and region (n = 1,103). **B)** Volcano plot of cortex vs nucleus sample proteins. Significant protein groups are shown in red with cut-offs of log_2_FC > 1.5 or log_2_FC < -1.5 and p value < 0.05 as calculated from Welch’s t-test.

Next, we validated the separation of the zebrafish lens cortex and nucleus regions observed proteins in the diaPASEF dataset. A PCA plot using untransformed MaxLFQ quantities for 1,103 protein groups found in all diaPASEF samples showed that the samples were separated into two distinct clusters: cortical and nuclear (Figure 4A). The same pattern was observed in the orbitrap DIA data (Supplemental Figure 5A).

Additionally, protein abundances were compared between the nuclear and cortical diaPASEF datasets via a volcano plot to identify differentially expressed proteins. No imputation was performed and only proteins found in all three replicates were considered (Figure 4B). The results revealed that several ribosomal proteins such as Rps8, Rpl8, Rpl13, and Zgc:171772 were differentially expressed in the cortex relative to the nucleus. The differential expression of ribosomal proteins is consistent with known biology in that protein translation occurs in cortical fiber cells and that organelles are degraded during fiber cell differentiation, leading to a loss of ribosomal proteins in fully differentiated nuclear fiber cells.^1^ Our results also show that Gstp1.2, an enzyme involved in the conjugation of glutathione, is differentially expressed in the cortex. Glutathione is a major lens antioxidant important in preventing cataract formation.^32,33^

In the lens nucleus, structural membrane proteins, such as Lim 2.5, Aqp0a, and Aqp0b were differentially expressed.^34^ In particular, Aqp0a, Aqp0b, Gja8b, and Slc20a1b are all part of the lens microcirculation system that imports nutrients and exports waste products to and from the lens nucleus.^35–38^ Metabolically active proteins, such as Enpp6 and Lctlb were also differentially expressed in the lens nucleus, suggesting that some metabolically active proteins are retained after lens fiber cell differentiation. Lctlb has also been shown to be involved in lens suture and cataract formation in mice.^39,40^ These results showed that overall, more lens-specific proteins expressed during lens fiber cell differentiation were found in the nuclear region, indicative of the successful separation of the lens cortex and nucleus. A full list of significant proteins, as visualized in the volcano plot, is provided in Supplemental Table 1. Similar trends were also observed in the orbitrap DIA method volcano plot (Supplemental Figure 5B). Significant proteins in the orbitrap DIA method volcano plot are shown in Supplemental Table 2. Only 5 significant proteins were found in both datasets, Rpl13, Gstp1.2, Si:dkey-164f24.2, Lctlb, and Aqp0b. Of the 24 remaining non-overlapping significant proteins identified in the DIA dataset, 23 were not significant in the diaPASEF dataset, and 1 was detected exclusively in the DIA dataset. However, 22 out of these 23 proteins met the *p* value cut-off but did not meet the log_2_FC cut-off. Therefore, in the diaPASEF dataset there was a statistically significant but small difference between the expression of these proteins in the cortex and the nucleus. This can be due to differences such as duty cycle, dynamic range, and sensitivity between the instruments and methods.

**Table 1.** Overrepresentation analysis of statistically significant cortical diaPASEF proteins (*p* value < 0.05 and log_2_FC > 1.5).

| GO Term | Annotation | # in Set | Fold Enrichment | FDR |
| --- | --- | --- | --- | --- |
| **GO Biological Process** |  |  |  |  |
| GO:0006412 | Translation | 15 | 7.94 | 8.55E-09 |
| **GO Molecular Function** |  |  |  |  |
| GO:0003735 | Structural constituent of ribosome | 15 | 16.4 | 0.344 |
| **GO Cellular Component** |  |  |  |  |
| GO:0022625 | Cytosolic large ribosomal subunit | 12 | 24.32 | 1.54E-13 |

Finally, we performed an overrepresentation analysis for significant proteins identified via Welch’s t-test for both diaPASEF cortical and nuclear datasets using PANTHER.^26^ In the cortex diaPASEF dataset, there is an overrepresentation of ribosomal and translation-associated gene ontologies (Table 1).

**Table 2.** Overrepresentation analysis of statistically significant nuclear diaPASEF proteins (*p* value < 0.05 and log_2_FC < -1.5).

| GO Term | Annotation | # in Set | Fold Enrichment | FDR |
| --- | --- | --- | --- | --- |
| **GO Biological Process** |  |  |  |  |
| No statistically significant results |  |  |  |  |
| **GO Molecular Function** |  |  |  |  |
| GO:0005372 | Water transmembrane transporter activity | 2 | > 100 | 5.20E-02 |
| GO:0004096 | Catalase activity | 2 | > 100 | 3.46E-02 |
| **GO Cellular Component** |  |  |  |  |
| GO:0005911 | Cell-cell junction | 4 | 11.36 | 2.31E-02 |
| GO:0005886 | Plasma membrane | 10 | 4.33 | 2.95E-03 |
| GO:0005737 | Cytoplasm | 3 | 0.28 | 8.07E-03 |

Conversely, the diaPASEF nucleus dataset exhibits an overrepresentation of water transmembrane transporter activity, catalase activity, cell-junction, cytoplasmic, and membrane gene ontologies (Table 2). These gene ontology results further validate the successful separation of the younger organelle-containing lens cortex from the older organelle-free lens nucleus. Overrepresentation analysis results for the orbitrap DIA cortex dataset are shown in Supplemental Table 3. No overrepresentation analysis was performed on the orbitrap DIA nucleus dataset, due to the low number of significantly expressed proteins.

## Conclusion

We have shown that a 30-minute liquid chromatography diaPASEF method on a timsTOF instrument identifies more peptide and protein groups in the lens than a 95-minute liquid chromatography DIA method on an orbitrap instrument due to a faster acquisition time and the addition of ion mobility. Thus, using diaPASEF, we have collected the most comprehensive zebrafish lens proteome dataset to date. Through the analysis of the diaPASEF dataset, we have also demonstrated the first instance of successful separation of the zebrafish lens cortex and nucleus regions. diaPASEF is a methodology that will allow further probing of the lens proteome using model organisms to more thoroughly understand lens biology.

## Supporting Information

Median normalization boxplots, peptide intensity percent CV per sample, venn diagrams comparing overlap between regions, venn diagrams comparing overlap between replicates, orbitrap DIA PCA and volcano plots, tables of significant proteins identified in diaPASEF and orbitrap DIA datasets, significant GO terms for orbitrap DIA cortex dataset (PDF)

## Acknowledgments

We thank Zhen Wang, Lee Cantrell, and the Mchaourab lab for critical discussions about this manuscript. The authors would also like to acknowledge support from the Vanderbilt Mass Spectrometry Research Center and NIH grants T32 GM065086 (SRZ), T32 EY007135 (SRZ), R01 EY012018 (HSM), P30 EY008126 (KLS), and R01 EY024258 (KLS).

