## Supplemental Figures 1-5, Supplemental Tables 1-3 for "Data-independent acquisition parallel accumulation-serial fragmentation (diaPASEF) analysis of the separated zebrafish lens improves identifications"

### Supplemental Materials

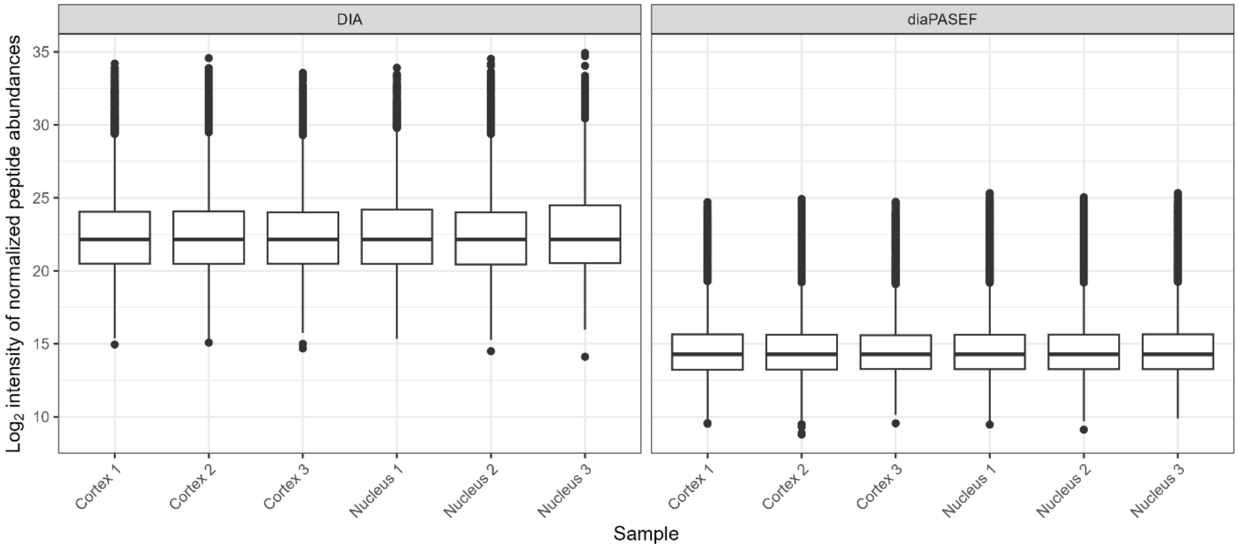


**Supplemental Figure 1. $Log_2$ quantity of peptide MS2 intensities after median normalization on DIA data collected on a Thermo Exploris 480 instrument and diaPASEF data collected on a Bruker timsTOF HT instrument.**

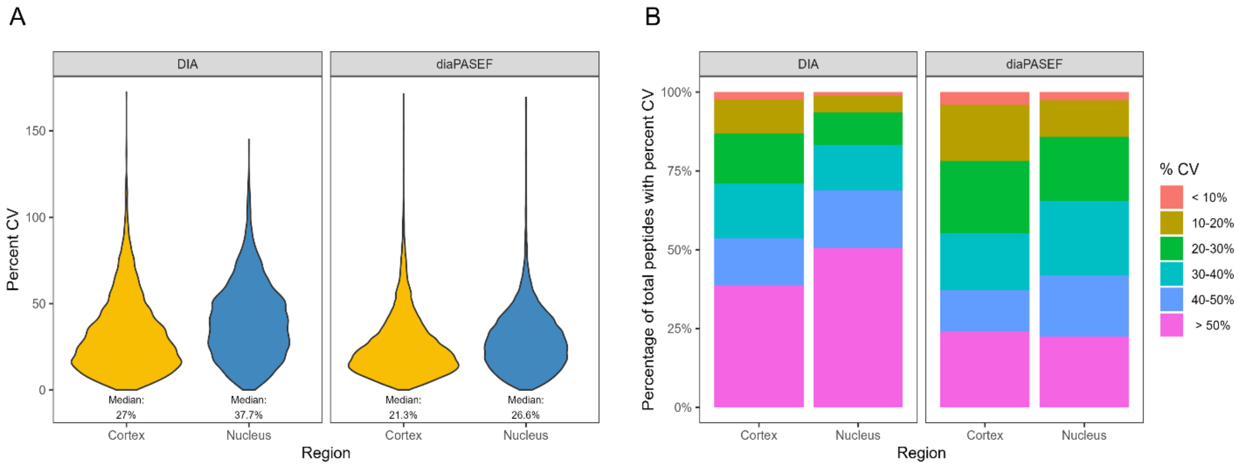


**Supplemental Figure 2. Peptide intensity % CV results for the separated zebrafish lens cortex and nucleus.** All peptides with < 1% q-value and at least 2 unique peptides per protein group were considered. Results were not separated by biological replicate or filtered to include only unique peptides. In total, 9,540 peptides were analyzed for the orbitrap DIA cortex samples, 8,087 peptides were analyzed for the orbitrap DIA nucleus samples, 40,211 peptides were analyzed for the diaPASEF cortex samples, and 33,492 peptides were analyzed for the diaPASEF nucleus samples. **A)** Violin plot of % CV for cortical and nuclear peptides for data collected using both DIA on a Thermo Exploris 480 and diaPASEF on a Bruker timsTOF HT. **B)** Bar plot of % CVs for cortical and nuclear peptides broken down by percentage of peptides within a specific range of % CV.

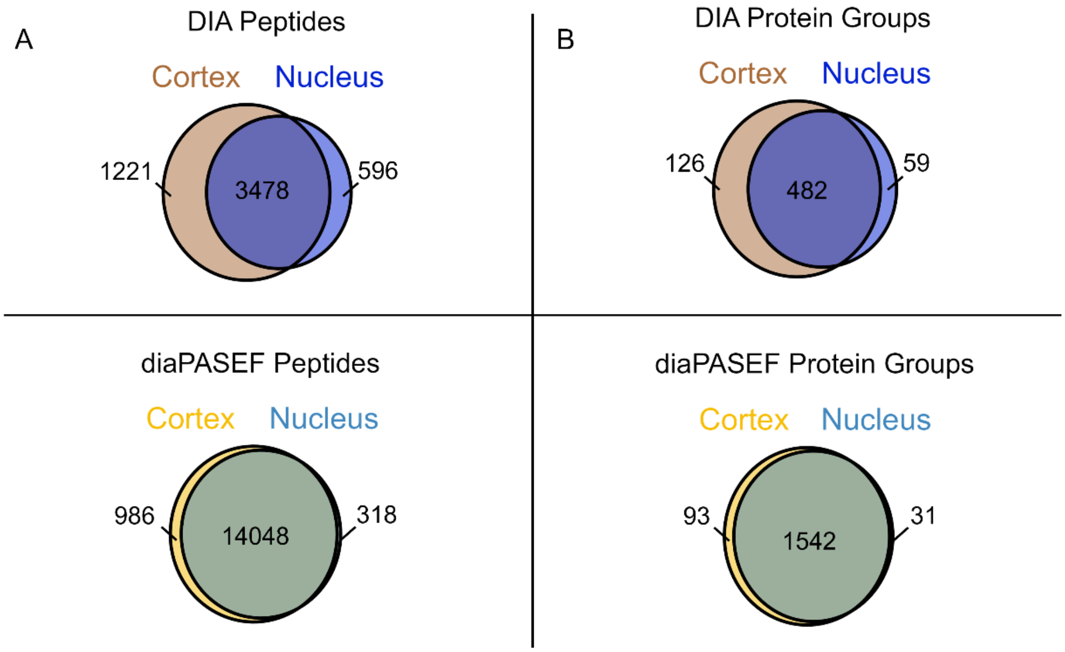


**Supplemental Figure 3. Venn diagrams showing the overlap between A)** peptides and **B)** protein groups between the DIA and diaPASEF data collected on the separated zebrafish lenses. All peptides and protein groups found in at least one biological replicate were compared, with duplicates removed.

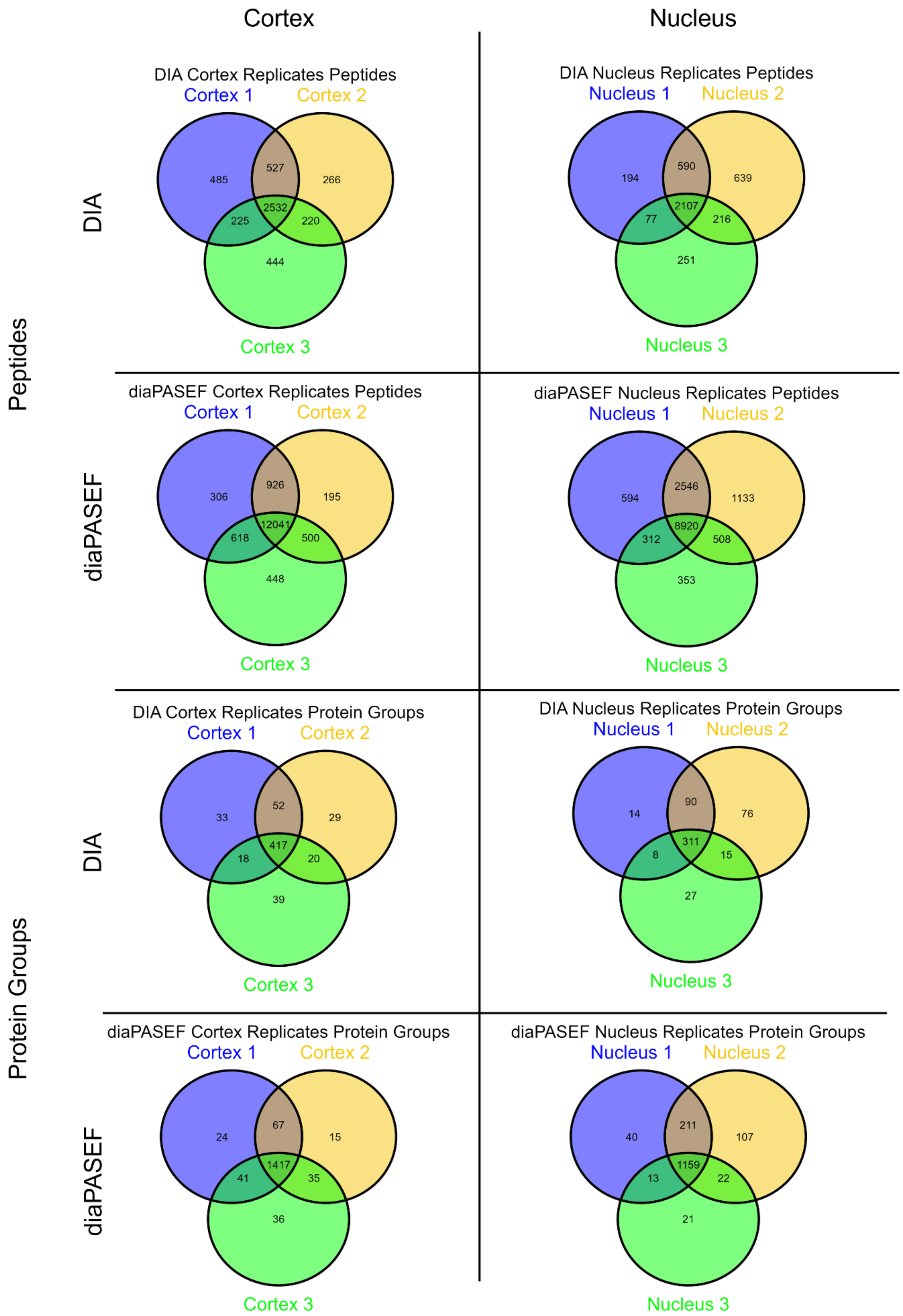


**Supplemental Figure 4. Venn diagram showing the overlap between three biological replicates for zebrafish lens cortex and nucleus samples on the peptide and protein group level separated by data acquisition method.**

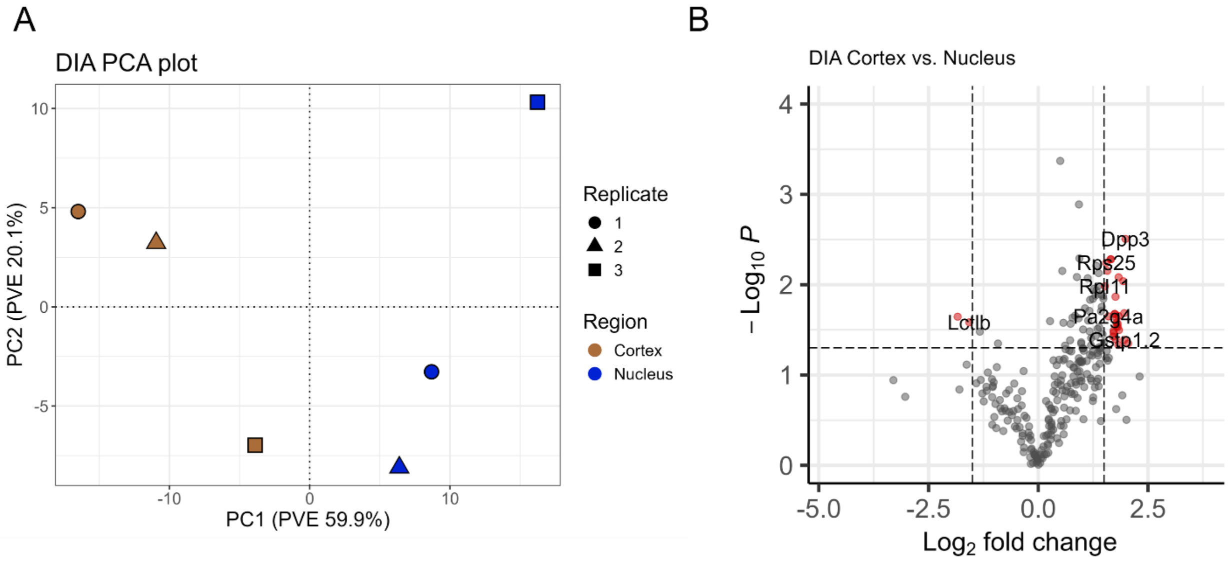


**Supplemental Figure 5. Analysis results of orbitrap DIA data. A)** PCA plot separated by replicate and region (n = 263). **B)** Volcano plot of cortex vs nucleus sample proteins. Significant protein groups are shown in red with cut-offs of $log_2FC > 1.5$ or $log_2FC < -1.5$ and p value < 0.05 as calculated from Welch's t-test.

**Supplemental Table 1. Significant proteins ($p$ value < 0.05 and $log_2FC$ > 1.5 or $log_2FC$ < -1.5) in the diaPASEF dataset organized by $Log_2FC$ (n = 35).**

| Gene Group | Protein Group | Protein Description | $p$ value | $Log_2FC$ |
|---|---|---|---|---|
| *rps8* | P62247 | Small ribosomal subunit protein eS8 | 0.003909041 | 2.481897098 |
| *rpl8* | Q6P0V6 | Large ribosomal subunit protein uL2 | 0.010171792 | 2.25857545 |
| *rpl27a* | Q6P2M0 | Large ribosomal subunit protein uL15 | 0.008776574 | 2.153349959 |
| *rpl36* | Q6Q415 | Large ribosomal subunit protein eL36 | 0.007524512 | 2.124129742 |
| *nat16* | E7FET0 | N-acetyltransferase 16 | 0.031498825 | 2.070295384 |
| *rpl34* | Q7ZWJ7 | Large ribosomal subunit protein eL34 | 0.003563581 | 2.069496022 |
| *rpl10* | Q7ZV96 | Large ribosomal subunit protein uL16 | 0.013188797 | 2.036976369 |
| *faua;faub* | F1RA98;Q6PC01 | FAU ubiquitin-like and ribosomal protein S30 fusion b | 0.006760121 | 2.019043612 |
| *rpl21* | Q6IQQ0 | 60S ribosomal protein L21 | 0.011212962 | 2.002765545 |
| *rbbp9* | Q1MT41 | Hydrolase RBBP9 | 0.006132191 | 1.925186015 |
| *rpl26* | Q7SXA1 | Large ribosomal subunit protein uL24 | 0.009230474 | 1.924398846 |
| *rpl13* | Q90Z10 | Large ribosomal subunit protein eL13 | 0.019915798 | 1.91269277 |
| *rps26l* | Q6PBZ2 | 40S ribosomal protein S26 | 0.011059105 | 1.898059165 |
| *rps24* | A0A8M2B343 | 40S ribosomal protein S24 | 0.012945126 | 1.77181796 |
| *rpl28* | F1QFV6 | Large ribosomal subunit protein eL28 | 0.008572089 | 1.767039669 |
| *rpl31* | Q24JV3 | Large ribosomal subunit protein eL31 | 0.007176367 | 1.765724655 |
| *zgc:171772* | A7YY10 | 60S ribosomal protein L37a | 0.013683676 | 1.709087001 |
| *rpl7a* | Q6PBZ1 | 60S ribosomal protein L7a | 0.000688221 | 1.638893239 |
| *gstp1.2* | Q9DDU5 | Glutathione S-transferase | 0.030139252 | 1.555262716 |
| *si:dkey-164f24.2* | A0A286Y8V3 | Si:dkey-164f24.2 | 0.018064922 | 1.533536117 |
| *cav1* | Q6YLH9 | Caveolin | 0.033681486 | -1.502378174 |
| *atp2b1a* | A0A8M9Q5T9 | Calcium-transporting ATPase | 0.005726974 | -1.521883362 |
| *lctlb* | A0A286Y8V0 | Lactase-like b | 0.033393737 | -1.606722692 |
| *slc20a1b* | Q6PFM1 | Sodium-dependent phosphate transporter 1-B | 0.015976032 | -1.632235297 |
| *tmem47* | Q6PFT6 | Transmembrane protein 47 | 0.013242156 | -1.665511898 |
| *rac1b* | Q29RC5 | Ras-related C3 botulinum toxin substrate 1 | 0.033802902 | -1.688233208 |
| *misp* | A0A0R4IHS7 | Mitotic interactor and substrate of PLK1 isoform X1 | 0.034788062 | -1.733434488 |
| *enpp6* | Q5BKW7 | Glycerophosphocholine cholinephosphodiesterase ENPP6 | 0.003057799 | -1.791046449 |
| *zmp:0000000625* | A0A8M1RIF4 | Cadherin-2-like | 0.021237841 | -1.833211042 |
| *si:dkey-57a22.15* | B0UYP0 | Gamma-crystallin A | 0.036082097 | -1.913493079 |
| *lim2.5* | Q5BKW4 | Lens intrinsic membrane protein 2.5 | 0.001665345 | -1.967881439 |
| *aqp0a* | Q6DEI6 | Aquaporin-0a | 0.009523115 | -2.040109434 |
| *gja8b* | Q503J6 | Gap junction protein | 0.024193848 | -2.058087606 |
| *aqp0b* | Q4ZJI3 | Aquaporin-0b | 0.006478904 | -2.239139865 |
| *lim2.3* | Q6DGY5 | Lens intrinsic membrane protein 2.3 | 0.021196919 | -2.782351794 |

**Supplemental Table 2. Significant proteins (*p* value < 0.05 and $log_2FC > 1.5$ or $log_2FC < -1.5$) in the orbitrap DIA dataset organized by $Log_2FC$ (n = 29).**

| Gene Group | Protein Group | Protein Description | *p* value | $Log_2FC$ |
|---|---|---|---|---|
| *glo1* | Q6P696 | Lactoylglutathione lyase | 0.044777548 | 2.046854321 |
| *dpp3* | Q6DI20 | Dipeptidyl peptidase 3 | 0.003115512 | 1.986749978 |
| *gstp1.2* | Q9DDU5 | Glutathione S-transferase | 0.04077937 | 1.960622523 |
| *si:dkey-164f24.2* | A0A286Y8V3 | Si:dkey-164f24.2 | 0.020644478 | 1.959728754 |
| *rpl13* | Q90Z10 | Large ribosomal subunit protein eL13 | 0.009156845 | 1.937545666 |
| *rpl24* | Q8JGR4 | Large ribosomal subunit protein eL24 | 0.031828165 | 1.838986254 |
| *rps5* | Q6PC80 | Small ribosomal subunit protein uS7 | 0.008268908 | 1.830822172 |
| *eif3a* | Q6PCR7 | Eukaryotic translation initiation factor 3 subunit A | 0.022164875 | 1.825577397 |
| *rpl23* | Q6PC14 | Large ribosomal subunit protein uL14 | 0.046451327 | 1.823766909 |
| *eno1a* | A0A2R8Q1X2 | Phosphopyruvate hydratase | 0.029408901 | 1.812812676 |
| *rpl14* | Q6DRN7 | Large ribosomal subunit protein eL14 | 0.027148089 | 1.810202976 |
| *rps4x* | Q642H9 | Small ribosomal subunit protein eS4 | 0.026320039 | 1.781929636 |
| *rps14* | Q6PBW3 | Small ribosomal subunit protein uS11 | 0.023545001 | 1.780290144 |
| *aldoaa* | Q803Q7 | Fructose-bisphosphate aldolase | 0.028904612 | 1.779777615 |
| *ppiab* | Q6PC53 | Peptidyl-prolyl cis-trans isomerase | 0.040207925 | 1.766863159 |
| *rps13* | Q6IMW6 | Small ribosomal subunit protein uS15 | 0.013603921 | 1.760065774 |
| *rps23* | A8KB78 | Small ribosomal subunit protein uS12 | 0.021091342 | 1.7441086 |
| *phgdh* | F1QEY8 | D-3-phosphoglycerate dehydrogenase | 0.021081516 | 1.738861491 |
| *hspb1* | A4VAK4 | Heat shock protein beta-1 | 0.033953265 | 1.716295077 |
| *rps18* | Q8JGS9 | Small ribosomal subunit protein uS13 | 0.032440526 | 1.714271327 |
| *mapk1* | Q7ZW72 | Mitogen-activated protein kinase | 0.03757024 | 1.705442809 |
| *ass1* | Q66I24 | Argininosuccinate synthase | 0.005244634 | 1.655897175 |
| *rpl7* | Q6Q417 | Large ribosomal subunit protein uL30 | 0.005221744 | 1.649584684 |
| *pa2g4a* | Q8AW82 | Novel protein similar to human proliferation-associated 2G4 protein (PA2G4) | 0.022440603 | 1.597685365 |
| *rps16* | Q1LWH1 | 40S ribosomal protein S16 | 0.007020197 | 1.572988422 |
| *rps25* | Q6PBI5 | Small ribosomal subunit protein eS25 | 0.005709555 | 1.549377184 |
| *rpl11* | Q6IQI6 | Large ribosomal subunit protein uL5 | 0.010395899 | 1.518867893 |
| *lctlb* | A0A286Y8V0 | Lactase-like b | 0.026071736 | -1.573217191 |
| *aqp0b* | Q4ZJI3 | Aquaporin-0b | 0.022614919 | -1.837531594 |

**Supplemental Table 3. Overrepresentation analysis of statistically significant cortical orbitrap DIA proteins ($p$ value < 0.05 and $log_2FC$ > 1.5).**

| GO Term | Annotation | # in Set | Fold Enrichment | FDR |
|---|---|---|---|---|
| **GO Biological Process** | | | | |
| GO:0042254 | Ribosome biogenesis | 6 | 11.19 | 6.22E-03 |
| GO:0006412 | Translation | 13 | 5.1 | 6.81E-04 |
| **GO Molecular Function** | | | | |
| GO:0019843 | rRNA binding | 5 | 23.68 | 4.12E-04 |
| GO:0003735 | Structural constituent of ribosome | 14 | 11.34 | 8.87E-10 |
| **GO Cellular Component** | | | | |
| GO:0022627 | Cytosolic small ribosomal subunit | 7 | 14.36 | 2.16E-05 |
| GO:0022625 | Cytosolic large ribosomal subunit | 6 | 9.01 | 1.96E-03 |
| GO:0016020 | Membrane | 0 | < 0.01 | 0.0302 |
